# Long-term effects of xenotransplantation of human enteric glia in immunocompetent rats with brain injury

**DOI:** 10.1101/2024.11.15.623633

**Authors:** Nina Colitti, Edwige Rice, Franck Desmoulin, Maylis Combeau, Mélissa Parny, Lorenne Robert, Etienne Buscail, Barbara Bournet, Nathalie Vergnolle, Isabelle Raymond-Letron, Isabelle Loubinoux, Carla Cirillo

## Abstract

**Background:** Acute brain injury is characterized by extensive tissue damage, resulting in neuronal loss and functional deficits in patients. The capacity of nerve tissue to self-regenerate is insufficient to repair damaged tissue, thus therapies based on exogenous cells are urgently needed. Human enteric glia (EG) have interesting intrinsic properties that make them a valuable candidate for regenerative medicine. In this long-term study, we investigated whether human EG treatment induces tissue repair and improves functional recovery in a rat model of brain injury.

**Methods:** Acute brain injury was induced by malonate injection in the motor cortex of female rats, causing extensive tissue damage and long-lasting sensorimotor deficits. Human EG were isolated from gut tissue, expanded and administered intranasally in awake immunocompetent rats. To determine the long-term safety and efficacy of human EG treatment, longitudinal evaluation of sensorimotor function, *post-mortem* tissue regeneration and the fate of human EG were assessed thirty-six weeks after intranasal administration.

**Results:** Transplanted human EG satisfied the safety criteria, non-immunogenic and non-tumorigenic, required for cell therapy; they were well tolerated in immunocompetent rats, and induced sensorimotor improvement. Importantly, thirty-six weeks post-treatment, intranasally delivered human EG were detected in the rat brain, mainly in the injured motor cortex. This indicated that transplanted human EG migrated and successfully engrafted and integrated with the host tissue. Additionally, human EG induced tissue regeneration by enhancing endogenous angiogenesis and neurogenesis. Notably, thirty-six weeks after administration, human EG generated mature neurons that were enveloped by oligodendrocytes and formed synaptic connections with the host tissue.

**Conclusions:** Transplanted human EG induced tissue repair and showed regenerative potential after brain injury. This is the first study demonstrating the feasibility, safety and efficacy of intranasal administration of human EG for treatment of brain injury.

## Background

Acute brain injuries are associated with significant morbidity and mortality worldwide[1]. Patients frequently develop long-term disabilities, including sensorimotor deficits, as a result of damage to brain tissue and extensive cell loss. Self-repair can occur through endogenous mechanisms, including neurogenesis, gliogenesis and angiogenesis, but these are dramatically limited, especially in adulthood[2].

In recent decades, the use of stem/progenitor cells as a therapeutic strategy for brain tissue repair has been extensively tested in animal models, and, in some cases, in clinical trials in patients[3–8]. Various cell sources have been investigated, including neural, embryonic, mesenchymal, and induced pluripotent stem cells[9–13]. While encouraging results have been obtained in animal models, the translation of these findings to clinical application has proven to be challenging for several reasons. These include difficulties in choosing an appropriate route of administration; concerns about the safety of transplanted cells; the clearance of donor cells and poor integration with the host tissue[14]. Despite considerable progress, the field has yet to identify an ideal donor cell population.

A suitable cell candidate should, in principle, be easily accessible, demonstrate high plasticity (for the generation of neurons) while simultaneously exhibiting a minimal risk to cause adverse outcomes. In addition, there is a need for a ‘ready-to-use’ cell source that does not require complex *in vitro* manipulation with high production costs.

The objective of this study is to overcome the intrinsic issues associated with current cell-based therapies, by proposing an innovative cell source to help tissue repair after brain injury.

In humans, nerve tissue is not only present in the brain; it is also present in the periphery. The enteric nervous system (ENS) is a highly organized network of neurons and glia cells in the gastrointestinal tract, and may serve as interesting reservoir of transplantable cells[15]. Particularly, enteric glia (EG) in the ENS have properties that make them compelling candidates for regenerative purposes. In humans, EG have important role in neuronal support and display high plasticity[16–23]. In previous studies, we have demonstrated that human EG can be accessed with relative ease from patient intestinal tissue and isolated to obtain pure primary cultures[16,19,23]. Importantly, in comparison to stem/progenitor cells, we have shown that human EG can be readily expanded *in vitro* and cryopreserved[16], thereby allowing the quantities required for repair therapy to be obtained.

In this study, we investigated whether human EG transplantation has beneficial effects on functional outcome and tissue repair in a rat model of acute brain injury causing long-lasting sensorimotor deficits[24–26]. The model consists in the injection of a toxin, malonate, into the motor cortex, resulting in extensive tissue damage due to the inhibition of a reversible succinate dehydrogenase. This produces secondary excitotoxic lesions similar to those of focal ischaemia-reperfusion, mimicking ischaemic stroke, as we previously validated[25,26,35].

In injured animals, invasive cell transplantation induces exacerbation of tissue damage[27–31]. Intranasal administration represents a viable alternative for the successful delivery of donor cells, which cross the cribriform plate and migrate *via* the rostral migratory stream to the brain[32–34]. Here, we demonstrate that, following intranasal administration, human EG migrated to, and successfully engrafted, the brain tissue, where they were able to induce tissue repair after brain injury. Thirty-six weeks after transplantation, we observed enhanced endogenous angiogenesis and neurogenesis, together with new-tissue formation. In addition, once in the brain, human EG generated mature neurons that were surrounded by oligodendrocytes and formed synapses with the host tissue.

## Methods

### Animals

The study involved 28 female Sprague-Dawley rats (280 to 320g, 11 weeks old, Janvier, France). They were housed in pairs (cage size: 30 cm in length, 18 cm height, 32 cm width) in a controlled environment (20°C) with a 12h/12h light/dark cycle and had free access to food and water. The protocol was approved by the “Direction Départementale de la Protection des *Populations de la Haute – Garonne*” and the “Comité d’éthique pour l’expérimentation animale *Midi-Pyrénées*” (protocol n° APAFIS#22419 2019101115259327v5, approved in 2019). Every effort was made to minimize the number of animals used and their distress. The animals were treated in accordance with the guidelines issued by the Council of the European Communities (EU Directive 2010/63). The work has been reported in line with the ARRIVE guidelines 2.0. The animals were divided into four groups: rats receiving human EG *(n=7)*; rats receiving vehicle (PBS, *n=7*); rats receiving human EG-conditioned medium *(n=7*, “conditioned medium” in the text) and rats receiving medium alone (DMEM, *n=7*). Based on our previous experiments and the statistical power obtained using PowerG software, the number of animals was calculated.

### Brain injury induction

To induce brain injury, we used the malonate model, based on the inhibition of a reversible succinate dehydrogenase, which produces secondary excitotoxic lesions similar to those of focal ischaemia-reperfusion[25,26,35]. This strategy is used to make an animal model with ischaemic brain injury[36–39]. Additionally, the malonate model is suitable for group comparisons in pharmacological studies involving time-consuming experiments (behaviour, MRI, histology). The rats were anesthetized with isoflurane (3% for induction, 3-5% for maintenance, in 0.7 litre/minute O_2_, Minerve^®^ compact anaesthesia workstation and monitoring, France) and restrained in a stereotaxic frame (Bioseb Laboratory, France). Cortical lesion of the motor area (M1) was induced by malonate injection [5 μL, 3M solution, pH 7.4 in phosphate buffer saline (PBS); Sigma-Aldrich, France] in *n=28* rats at the following stereotaxic coordinates: 2.5 mm lateral and 0.5 mm ahead to Bregma, with 2 mm depth[40]. The lesioned hemisphere was identified as the dominant one through the utilization of the grip strength test. This model was previously validated and published by our group[25,26]. Animals received general anaesthesia (isoflurane 3%) with analgesia (buprenorphine, 25 μg/kg) and methylprednisolone (20 mg/kg), to limit pain prior to surgery and control the oedema consequent to the brain injury. Animals were monitored once daily for the week following surgery and received post-operative analgesia (at 6 hours). DietGel^®^ was administered 24 hours prior to surgery and after surgery to facilitate feeding and hydration to aid the animal’s recovery. Given the duration of the experiment (up to 36 weeks), a multiple enrichment system was established. An enrichment object (tunnel, rope, coloured balls) was changed every month to meet the needs of the species. As part of the implementation of refinement strategies, we trained the animals to receive intranasal instillations without being anesthetized and/or restrained. This avoided repeated stress caused by anaesthesia. Signs of distress observed during daily observation, such as isolation and indifference to the external environment, a reduction in exploratory behaviour, a prone position with hunched back, flight or resistance to handling, or weight loss >20% (weighed twice a week, then daily in the case of weight loss) were established as humane endpoints. 48 hours and 72 hours after surgery no animal showed the above listed signs or weight loss >20%.

### Behavioural tests

The study was conducted during the Coronavirus pandemic, with a restricted presence of zootechnicians and experimenters during the behavioural testing phase. Notably, the emergence test demonstrated an increase in the level of anxiety in some rats (irrespective of the experimental groups) eight weeks after the injury, which corresponds to the commencement of the lockdown due to the Coronavirus pandemic. Accordingly, these values were excluded from the analysis of all behavioural tests, given that they may have been influenced by the animal’s anxiety status.

To assess sensorimotor function, rats were trained to perform a grip strength test and neurologic severity score two weeks before surgery. Behavioural tests were performed one week before injury (baseline), then 48hrs after injury, weekly for the first month, and monthly for the following six months. Each time point was assessed in triplicate on three different days within the same week. The limb-use asymmetry test and the light-dark box test were two additional tests used in the study. These were performed eight times: one week pre-injury, and then one-, four-, eight-, twelve-, sixteen-, twenty-, and twenty-four-weeks post-injury. For all tests, each value reported represents the median ± interquartile range [first quartile (Q1); third quartile (Q3)] of each group. The tests were conducted using the same protocol as a previous team study[24]. The animals were matched for grip and neurological scale performance, which correlates with lesion size, as shown in a previous study by the team[24] and assigned to the different groups (Vehicle vs. human EG; medium vs. conditioned medium). No animals were excluded from the study.

#### Emergence Test

The light-dark box test evaluates anxiety based on the aversion to bright light and spontaneous exploratory behaviour in a new environment. The latency to exit the dark side and enter the light compartment was recorded.

#### Grip Strength Test

The grip strength test measures the maximal forelimb muscle grip strength. The experimenter restricts the rat by holding it by the back and leaving the forelimbs free. Maximum isometric force (in Newtons) is measured on the attached dynamometer. This test determines the dominant paw of the rat. This test was validated in previous studies by our team[25,41]. Values of the contralateral forelimb were normalized by values of the ipsilateral forelimb.

#### Neurological Severity Scale (NSS)

The NSS includes five tests to evaluate sensorimotor function (reflexes, stability), sensitivity (proprioception) and depression. All the tests were scored in a scale from 0 to 16 points. The higher is the score, the more severe are the deficits.

#### Limb-Use Asymmetry Test

The asymmetry test allows the analysis of motor deficits of the whole limb (shoulder, elbow, paw) over time. Spontaneous use of the forelimb is evaluated in this test. The percentage of forelimb use was calculated as follows: Asymmetry (%) = number of times the ipsilateral paw is used for support - number of supports for the contralateral paw/total number of supports x 100.

### *In vivo* Magnetic Resonance Imaging

A 7T Biospec animal imager (Bruker Biospin, Ettlingen, Germany) was used for *in vivo* MRI. Rats were scanned at two time points: twelve- and twenty-four-weeks post-injury (*n*=7 per group). Rats were anaesthetized with isoflurane (3% for induction, 1% for maintenance, in 0.7 liter/minute oxygen, Minerve^®^ compact anaesthesia workstation and monitoring, France) prior to image acquisition. To maintain a body temperature of 37°C, the animals were placed in a thermoregulated imaging cell. To identify the lesion size and its evolution, anatomical imaging in T2 weighting using a T2 Turbo-RARE sequence with TE/TR = 35.7/5452 ms, Rare Factor = 8 and coronal slices with a spatial resolution of 0.137x0.137x0.500 mm3 were acquired. The acquisition time was 17 min 27 sec.

### Isolation, propagation, and cryopreservation of human EG

Human EG were isolated and propagated as we previously established[16]. Human gut tissue was obtained from n=3 patients (2 males and 1 female, age 68±8.7) undergoing surgery for colon resection, admitted to the Digestive Diseases Unit of the Centre Hospitalier Universitaire de Purpan (Toulouse). The biocollection was subject to an ethical protocol approved by the national application for the management of the conservation of elements of the human body (CODECOH, protocol: DC - 2015-2443). Informed consent was obtained from all subjects involved in the study. Human EG were isolated as we previously described[16]. In detail, after removal, the sample was placed in a chilled solution of sterile Krebs-Henseleit oxygenated solution, equilibrated to pH 7.4. The tissue was incised along the mesenteric border, laid flat and then placed with the mucosal layer up in a Petri dish containing Krebs-Henseleit solution, which was changed every five minutes. The mucosa and submucosa were removed under a dissecting microscope to reveal the myenteric plexus. This was then cut into thin sections. Subsequently, an enzymatic digestion was performed (45 minutes at 37°C with 1mg/mL protease, 1.25 mg/mL collagenase, and 1% sterile Bovine Serum Albumin, all from Sigma-Aldrich). The digestion was initiated by mechanical stirring. Isolated ganglia (containing neurons and EG) were selected under a dissecting microscope, and then placed in a culture dish covered with glial culture medium [Dulbecco’s modified Eagle’s medium (DMEM) - F12 mixture 1:1, supplemented with 10% inactivated fetal calf serum (FCS), and 1% antibiotic-antimycotic solution, all from Gibco)]. Cultures were kept at 37°C, 5% CO_2_ and 95% humidity. The confluence was reached in 2-3 weeks[16]. As previously established[16,19], this protocol allows to obtain pure cultures of human EG, which are devoid of “contaminating cells”, i.e. neurons, fibroblasts or smooth muscle cells. Cultures were expanded until the third passage. They were then trypsinized, transferred in a cryoprotectant medium (DMEM-F12 1:1 mixture, DMSO 10%) and transferred to liquid nitrogen[16,42]. The cryovials were thawed two weeks prior to the intranasal delivery in rats and passaged two times to obtain the required number of cells to administer. To ensure consistency, human EG were used at their fifth passage throughout the study. This protocol enabled the desired number of cells (1×10) to be obtained for transplantation.

### Characterization of human EG cultures

Characterization of human EG cultures was performed by immunofluorescence. To this aim, the cells were placed in an 8-compartment LAB-TEK^TM^ slide culture chamber. They were washed 3 x 10 minutes with PBS 1X before fixation in paraformaldehyde (PFA) 4% for 30 minutes at room temperature. PBS-Triton-X at 0.1 mol/L combined with 4% goat serum (blocking buffer) was used to mask non-specific binding sites for 2hrs at room temperature. Cells were then incubated overnight at 4°C with primary antibodies for glia, neurons and fibroblasts (Table 1). The cultures were then incubated for 2hrs at room temperature with respective secondary antibodies (Table1). The absence of non-specific labelling was ensured by negative controls.

**TABLE 1.** List of primary and secondary antibodies used for staining of hEG cultures.

| Primary antibodies (catalog #) | Host species | Dilution | Secondary antibodies |
| --- | --- | --- | --- |
| GFAP (Dako #Z0334) | rabbit | 1:1500 | Goat anti-rabbit 488 Alexa Fluor |
| S100 (Dako #Z0311) | rabbit | 1:500 | Goat anti-rabbit 568 Alexa Fluor |
| NeuN (Abcam #ab177487) | rabbit | 1:300 | Donkey anti-rabbit 594 Alexa Fluor |
| Alpha-SMA (Abcam #ab7817) | mouse | 1:500 | Donkey anti-mouse 594 Alexa Fluor |
| HuC/HuD (Abcam #ab191181) | mouse | 1:500 | Donkey anti-mouse 488 Alexa Fluor |
| ASCL1 (Mash1, Abcam #ab211327) | rabbit | 1:500 | Donkey anti-rabbit 594 Alexa Fluor |
| SOX10 (Santa Cruz Biotechnologies #sc-17342) | goat | 1:300 | Donkey anti-goat 488 Alexa Fluor |

### Human EG-conditioned medium preparation

To obtain the human EG-conditioned medium, Once the cells had reached confluence, they were washed with sterile PBS 1X and subsequently incubated in FCS-free medium (DMEM : F12 1:1 mixture). Following a 48-hour incubation period, the medium was collected, centrifuged at 1000g for 10 minutes, filtered through a 0.22 μm filter, and stored at −80°C in 500 μl aliquots. Prior to utilization, the medium was thawed by means of a water bath maintained at 37°C.

### Intranasal administration of human EG in immunocompetent rats

Ten days after brain injury, rats received human EG *(n=7)* or vehicle (PBS, *n=7*), *via* the olfactory pathway[43,44] using a 20 µl pipette with cone. Cell suspension was prepared as follows: 48hrs before intranasal delivery, human EG (passage fifth) were cultured in foetal calf serum (FCS)-free medium (DMEM: F12 1:1 mixture). To avoid problems in interpreting the data from the administration of human EG from three patients, we decided to use human EG from one patient. The day of intranasal administration, cells were washed with sterile PBS 1X, trypsinised (Trypsin-EDTA solution 1X, Sigma-Aldrich), centrifuged at 1400 rpm for 5 min. Next, the supernatant was discarded and the cells were resuspended at a density of 2.5×10^5^cells/60 μl in sterile PBS 1X. Thirty minutes prior to the administration of human EG, all animals were given 100 U hyaluronidase (Sigma-Aldrich) dissolved in sterile PBS 1X. Hyaluronidase was used to disrupt the cells tight junction and the barrier function of the nasopharyngeal mucosa to facilitate the entry of the cells into the brain[34,45,46]. Rats were awake and held under the forelimb by an experimenter. Intranasal delivery was performed once a week for one month. Ten μl drops containing either human EG suspension or vehicle were placed at the edge of one nostril and instilled (10 μl per nostril, three times, 60 µl/week). At the end of the treatment, a total volume of 240 μl of cell suspension (1×10^6^ cells) or vehicle was used, per animal. The volume and cell numbers were optimized and chosen based on the literature, to be effective in delivering cells into the brain through the cribriform plate[34,47–49]. In parallel, a third and fourth group of rats received conditioned medium *(n=7)* or medium alone (DMEM, *n=7*), respectively. In these groups, intranasal application was performed every day for one month (10 μl per nostril, three times, 60 µl/day). At the end of the treatment, a total volume of 600 μl of conditioned medium, or medium alone, were used. The experimenters were blinded to treatment throughout the whole study.

### Histological analysis of brain slices

The brain tissue was processed to evaluate tissue anatomy, remodelling, and new-tissue formation thirty-six weeks post-injury, which corresponds to thirty-two weeks after human EG intranasal administration. Nissl staining was used to evaluate tissue regeneration, while staining with specific markers was performed to identify human donor and host cells.

Rats were anesthetized with isoflurane and sacrificed with a lethal intraperitoneal injection of pentobarbital (160 mg/kg, Centravet). Intracardiac perfusion of heparinized 0.9% NaCl (200 ml, 20 min) eliminated blood from the vessels, and was followed by PFA 4% (250 to 300 ml, 40 min) for fixation. The brain was then extracted and immersed in a PFA 4% bath for 24h at 4°C and washed in two successive baths of PBS 1X. Sucrose baths of increasing concentration (10, 20 and 30%) were used for cryoprotection. Approximately 500 coronal brain slices (thickness: 20 μm) were cut using a microtome (Sliding Microtome Microm HM 450, Thermo Scientific, Germany) (*n=4* rats/group). Ethanol baths were used for *n=2* brains (one for human EG group and one for PBS group) to perform paraffin blocks.

#### Nissl and Ki67 staining

One in every twelve sections was stained with Nissl staining (Cresyl violet dye). Stained slides were scanned on a 3DHISTECH’s Slide Converter, and images used for the quantification of reconstructed tissue area. Quantification was performed following the method previously reported[24,50]. Ki67 immunohistochemical staining with DAB as chromogen was realized on 8 μm coronal brain slices from paraffin block. This marker is used to identify proliferating cells. Four slices per rat were used for the quantification of proliferating cells.

#### Immunostaining

Floating sections were incubated in PBS buffer 0.1% Triton X-100 (Sigma-Aldrich) and 4% serum (donkey or goat, Thermo Fisher Scientific or Sigma-Aldrich, respectively), for 2hrs at room temperature. Sections were exposed to the primary antibodies (Table 2), overnight at 4°C. After three washes with PBS 1X, sections were incubated with a specific secondary antibody coupled to a fluorochrome (Table 2), for 2hrs at room temperature. Images were captured using a Nikon Eclipse Ti2 series fluorescence microscope (Nikon Europe B.V., The Netherlands) or a confocal fluorescence microscope Zeiss LSM710 (Cell Imaging facility, Toulouse Institute for Infectious and Inflammatory Diseases, Inserm) and analysed using Fiji[51] and Imaris image analysis software. To evaluate tissue remodelling, gliosis, angiogenesis and neurogenesis were quantified in the reconstructed tissue in the injured region of the brain. Surface quantification was used for cytoplasmic markers: Gfap, Iba1, Doublecortin (Dcx) and beta 3 tubulin [three counting frames (645 x 628 μm^2^) per section, 20X magnification, *n=4* rats per group]. Based on the Nissl staining, adjacent sections were chosen, where the reconstructed tissue was identified. Mature neurons were quantified by counting their Neuron specific nuclear protein (NeuN)- positive nuclei in the reconstructed tissue [three counting frames per section, expressed as percent of DAPI^+^ cells, 20X magnification, *n=4* rats per group]. Blood vessels were quantified using the lectin-antibody [frames normalized by surface area (645 x 628 μm^2^ per section), 20X magnification, *n=4* rats per group]. We measured the glial barrier thickness following the method previously established by our team[24,50]. Human donor cells were identified and counted on entire coronal sections, acquired at 20X magnification, using the Addons Scanning Wizard function of the Nikon NIS-Element fluorescence microscope. The sections from 3.5mm bregma to −3mm bregma were analysed. To evaluate the location of human donor cells, the slices were divided into four groups (3.5 to 2mm bregma; 2 to 0mm bregma; 0 to −1.4mm bregma; −1.4 to −3mm bregma: twelve slices obtained from four rats per location group). A total of 48 brain sections were quantified. Identification of double markers for human cells were performed on four counting frames (645 x 628 μm^2^) for two sections per rat receiving human EG corresponding to *n=4*. A total of 32 fields were quantified.

**TABLE 2.** List of primary and secondary antibodies used for staining of rat brain slices.

| Primary antibodies (catalog #) | Host species | Dilution | Secondary antibodies |
| --- | --- | --- | --- |
| **GFAP (Dako #Z0334)** | rabbit | 1:1500 | Goat anti-rabbit 488 Alexa Fluor |
| **S100 (Dako #Z0311)** | rabbit | 1:500 | Goat anti-rabbit 568 Alexa Fluor |
| **Iba1 (Abcam #ab5076)** | goat | 1:500 | Donkey anti-goat 568 Alexa Fluor |
| **Lectin (Sigma L0401)** | Lycopersicon<br>esculentum | 1:200 | Conjugate with FITC |
| **DCX (SantaCruz #sc-8066)** | goat | 1:250 | Donkey anti-goat 568 Alexa Fluor |
| **βIII tubulin (Covance #PRB435P)** | rabbit | 1:500 | Donkey anti-rabbit 594 Alexa Fluor |
| **NeuN (Abcam #ab177487)** | rabbit | 1:300 | Donkey anti-rabbit<br>594 Alexa Fluor |
| **Human Nuclei (Abcam #ab254080)** | mouse | 1:100 | Donkey anti-mouse<br>594 Alexa Fluor |
| **STEM121 (TakaraBio Y40410)** | mouse | 1:400 | Donkey anti-mouse<br>594 Alexa Fluor |
| **OSP (Abcam #ab53041)** | rabbit | 1:200 | Goat anti-rabbit 488<br>Alexa Fluor |
| **Synaptophysin (Proteintech #17785-1-AP)** | rabbit | 1:500 | Goat anti-rabbit 488<br>Alexa Fluor |

### Statistical Analysis

Comparisons between two groups were performed using the Mann-Whitney U-test. Paired repeated measures were analysed using the mixed-effects analysis multiple comparison with Bonferroni correction. We used an unpaired t-test with a 95% confidence interval, after checking (with residual analysis) for equal variances between the different groups. Results were presented as median and the interquartile range (first quartile Q1; third quartile Q3). GraphPad Prism software was used for statistical analysis. Investigators were blinded to the treatment throughout the study.

## Results

The study protocol is shown in *Figure 1*. We had four groups of rats (human EG, vehicle, human EG-conditioned medium, and medium alone), in which the animal’s behaviour, brain anatomy (*in vivo* analysis), tissue regeneration and remodelling in brain slices (*post-mortem* analysis) were investigated. The strategy of using human EG-conditioned medium or medium alone allowed for the exclusion of potential effects due to factors other than cell delivery. By reason of clarity, it should be noted that the temporal discrepancy between the final MRI and the sacrifice was due to the constraints imposed by the Coronavirus pandemic, which necessitated a modification of the original plan.

**Figure 1.**
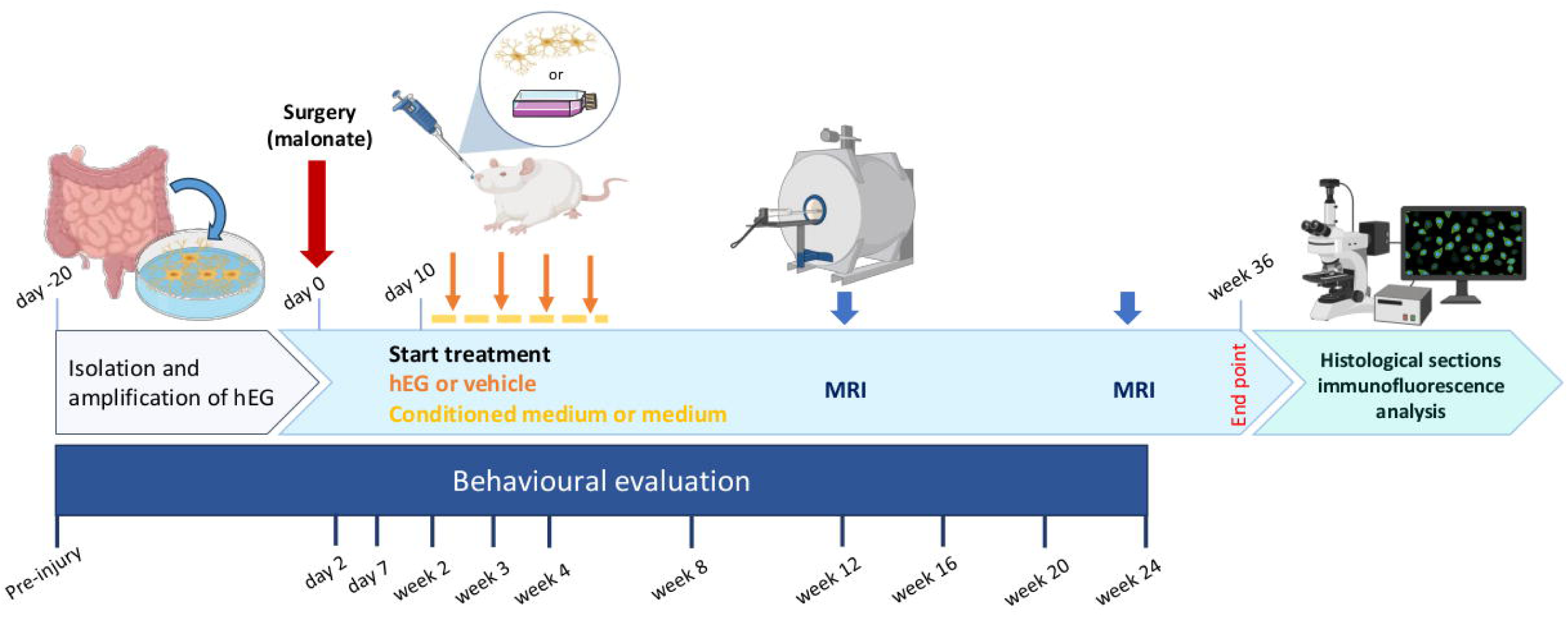
Detailed study protocol. Behavioural tests started before injury and were performed at different time points: day 2, day 7 and week 2, 3, 4, 8, 12, 16, 20 and 24 post-injury. Brain injury was induced in adult female rats (n=28, 12-week-old) by malonate injection into the cortex (red arrow). Ten days after the injury, rats received hEG (n=7) or vehicle (n=7) intranasally, once/week for a month (orange arrows; 2.5x10 cells/administration). The two other groups of rats received conditioned medium (n=7) or medium (n=7) ten days after injury every day for a month (yellow dashes). MRI acquisitions were performed 12 weeks and 24 weeks post-injury. Histological analysis was performed 36 weeks post-injury, to assess tissue regeneration, host cell reorganization, and to identify cell populations. MRI: magnetic resonance imaging. hEG: human enteric glia.

### *In vitro* characterization of the purity of human EG cultures

Mature human EG were isolated from the myenteric plexus of the gut and amplified as we previously established[16,19,23] (detailed in *Supplementary Methods)*. After five passages, before the intranasal administration in rats, human EG cultures were characterized by immunostaining to assess their purity. We evaluated the protein expression of glial specific markers: glia fibrillary acidic protein (GFAP), S100, SRY-Box Transcription Factor 10 (SOX10), which were all expressed in mature human EG (*Figure 2A-C*). As expected, primary cultures did not express fibroblast (alpha smooth muscle actin: α-SMA) (*Figure 2C, middle*) or neuronal markers (HuC/D and Neuronal nuclei: NeuN) (*Figure 2D-E*). These results demonstrated the purity of human EG cultures and the absence of contaminating cells, like neurons or fibroblasts, confirming our previous studies[16,19,23]. In addition, we investigated whether mature human EG in culture possessed neurogenic property. The expression of the transcription factor Anti-Achaete-scute homolog 1 (ASCL1), which regulates enteric neurogenesis[52], demonstrated that mature human EG have neurogenic characteristic (*Figure 2F*) and indicated that mature human EG do not need *in vitro* reprogramming to acquire this potential.

**Figure 2.**
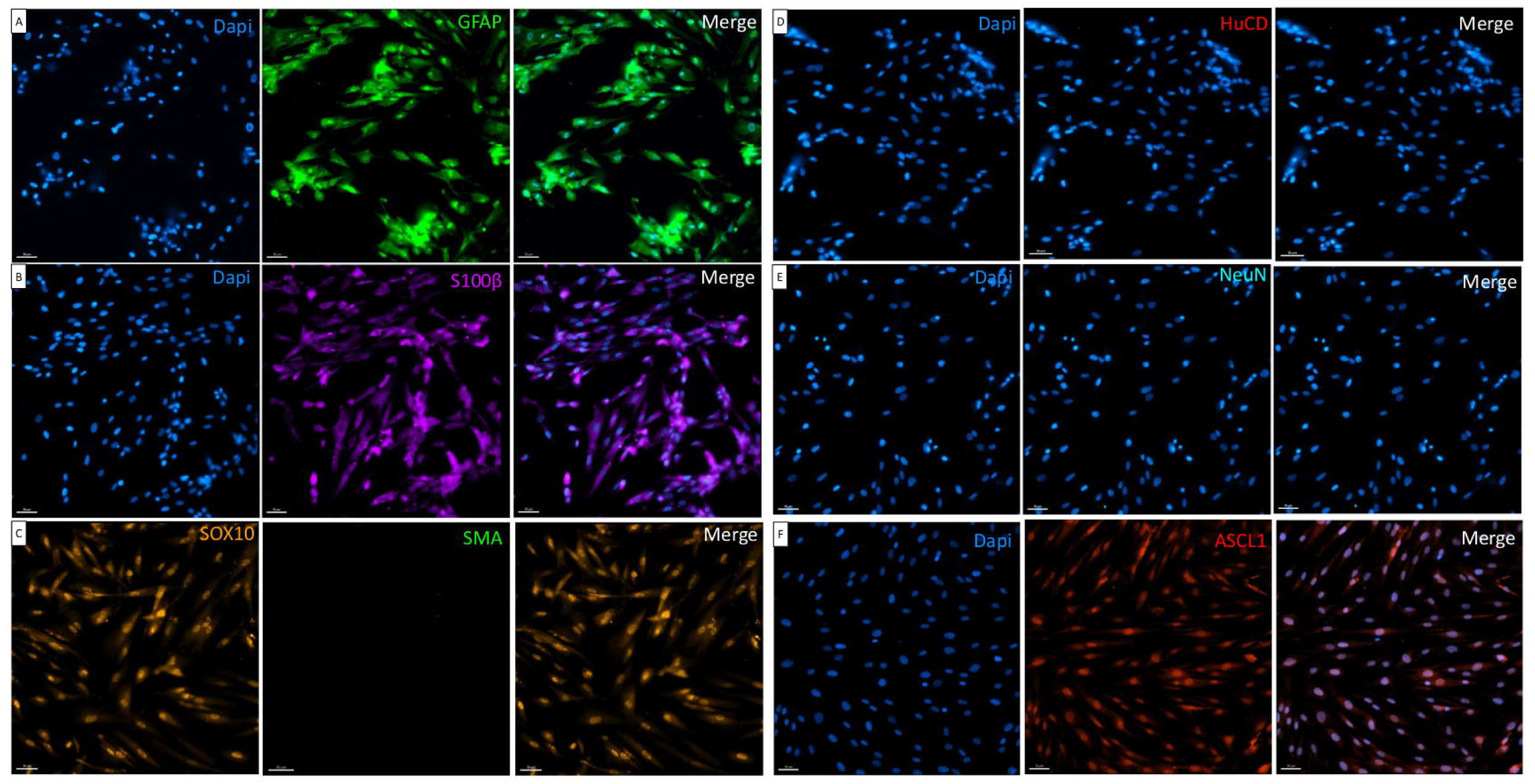
Immunostaining of hEG cultures. **(A)** Images showing Dapi labelling (left), GFAP positive labelling (middle) and merge (right). **(B)** Immunostaining of hEG showing Dapi labelling (left), S100ß positive labelling (middle) and merge (right). **(C)** Immunostaining of hEG showing positive nuclear SOX10 labelling (left), negative SMA labelling (middle) and merge (right). **(D)** Immunostaining of hEG showing Dapi labelling (left), negative HuCD labelling (middle) and merge (right). **(E)** Immunostaining of hEG showing Dapi labelling (left), negative NeuN labelling (middle) and merge (right). **(F)** Immunostaining of hEG showing Dapi labelling (left), positive nuclear ASCL1 labelling (middle) and merge (right). Scale bars: 50 μm. GFAP: Glial Fibrillary Acid Protein, S100ß: S100 intracytoplasmic calcium binding protein, SMA: alpha skeletal muscle, HuCD: neuron-specific RNA-binding protein, NeuN: Neuronal nuclei marker, ASCL1: Achaete-Scute Family BHLH Transcription Factor 1.

### *In vivo* characterization of brain injury after intranasal administration of human EG

MRI was performed twelve- and twenty-four weeks post-injury (*Supplementary* Fig. 1A and B). T2-weighted images allowed to visualize hyperintense, vasogenic and cytotoxic oedemas. Lesions are characterized by an increase in T2 signal. Twelve weeks post-injury, the distribution of lesion volumes in matched groups have similar medians: vehicle=79.15mm^3^ *vs* human EG=69.86mm^3^; medium=42.87mm^3^ *vs* conditioned medium=43.92mm^3^ (*Supplementary* Fig. 1C). The same tendency was measured twenty-four weeks post-injury, where the distribution of lesion volumes in matched groups have similar medians: vehicle=121.10mm^3^ *vs* human EG=94.15mm^3^; medium=55.50mm^3^ *vs* conditioned medium=51.72mm^3^ (*Supplementary* Fig. 1D). Regarding the distribution of atrophy in percentage, twenty-four weeks post-injury, the human EG group showed a reduced percentage, although not statistically significant, compared with the other groups: vehicle=6.20%; human EG=3.35%; medium=12.02%; conditioned medium=15.14% (*Supplementary* Fig. 1E).

### Transplantation of human EG shows long-term safety and induces sensorimotor improvement in immunocompetent rats

The observation of clinical signs provided regular monitoring of the animals after brain injury and intranasal administration throughout the study. The animals receiving human EG showed no signs of local or systemic toxicity, indicating that they tolerated well the treatment with exogenous human cells (xenotransplantation). This is an important finding given that our rats were immunocompetent. To further assess the safety of our therapeutic strategy, we used appropriate behavioural tests.

The emergence test provides information on anxiety state. The injection of malonate in the motor cortex increased the latency to exit the dark compartment one-week post-injury in all rats *(P<0.0001, Figure 3A*). As expected, the latency time was markedly diminished in all groups eight-week post-injury *(P<0.05*, *Figure 3A*), continuing until the twenty-four-week interval.

**Figure 3.**
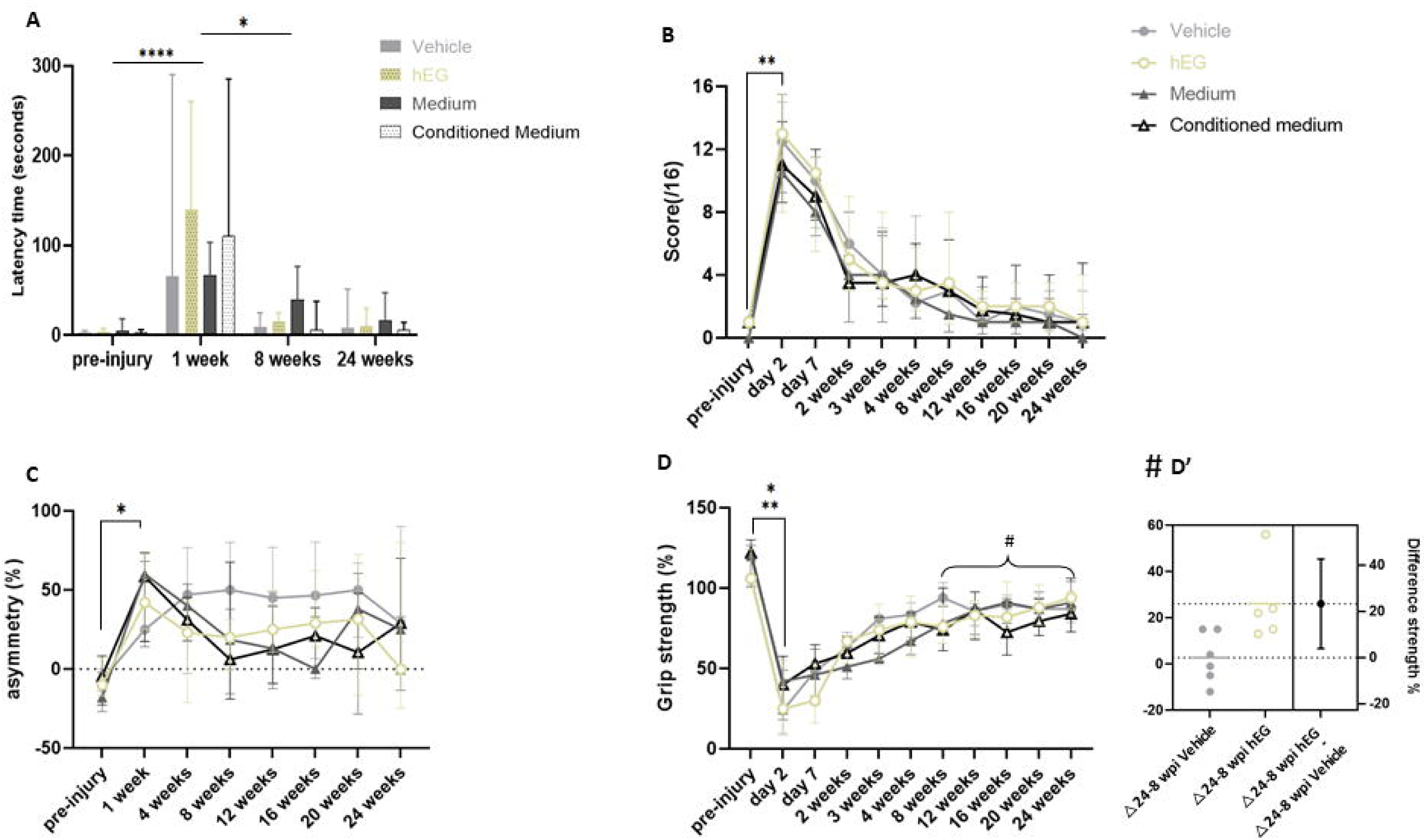
Effect of intranasal administration on behavioural tests. **(A)** The anxiety test measuring the latency to remove two paws from a black box. The latency was significantly increased one-week post-injury for all the groups (mixed-effects analysis multiple comparison test, *P<0.0001*). The latency significantly decreased between week and 8 weeks post-injury in all groups, independently of the treatment (*P<0.05*). **(B)** The neurologic scale score measures sensory-motor impairments (score out of 16). The four groups were significantly impaired 2 days post-injury (mixed-effects analysis multiple comparison test *P<0.01*). **(C)** The limb-use asymmetry test measures the asymmetric use of the limb. The four groups were significantly impaired on the contralateral limb one-week post-injury (mixed-effects analysis multiple comparison test *P<0.05*). **(D)** The grip strength test shows the grip strength of the front paw contralateral to the injected hemisphere compared to the other paw, expressed as %. Rats in all four groups were deficient two days after injury (Wilcoxon test, *P<0.05*). **(D’)** The graph shows the 95% confidence interval: [3.943 to 42.72] with a difference strength of 26% for hEG group and 2.6% for vehicle group between 8 weeks and 24 weeks. For (A–D): n=7 per group; wpi: weeks post-injury. The median and the interquartile range are represented.

The neurological severity scale (NSS) assessed the sensorimotor deficit measured with a scoring ranging from 0 to 16. *Figure 3B* shows a significant increase (*P<0.01*) in the score two days post-injury, reflecting the consequences of the lesion in the sensorimotor cortex. The rats recovered from twelve weeks post-injury onwards. A *plateau* was reached at twelve weeks. Statistically, there was no difference among the four groups.

The limb-use asymmetry test evaluated the spontaneous use of the front limbs. It measures the deficit of the whole limb including shoulder, elbow, and paw. The asymmetry corresponds to the difference between the use of the ipsilateral - contralateral paw compared to the number of supports, in percentage. Non-injured rats made simultaneous elevation against a support or used their dominant paw. As expected, the percentage of asymmetry increased in all groups one-week post-injury (*P<0.05, Figure 3C*). There was no significant effect among the four groups.

The grip test measures the grip strength of the forelimbs and was used to identify the dominant paw in each animal. The percentage of contralateral paw strength is shown in *Figure 3D*, after injury and after intranasal delivery of treatments. Rats in all four groups were deficient two days after injury. The mixed-effects analysis multiple comparison test compares the values measured two days post-injury to the pre-injury for the vehicle group (*P=0.012*), human EG group (*P=0.001*), medium group (*P=0.034*) and conditioned medium group (*P=0.006*). The recovery for vehicle and medium groups flattened to the top after spontaneous recovery (*Figure 2D*), as previously reported[24,25]. Rats receiving conditioned medium showed functional recovery comparable with the group receiving medium alone. Interestingly, the rats receiving human EG showed a good improvement in strength compared with vehicle group, with a better delta recovery calculated between eight and twenty-four weeks, *(t(9)= 2.72, p=0.023; IC_95_= [3.943- 42.72], Figure 3D’)*.

Together, behavioural tests indicated that all groups tolerated the treatments well, showing no worsening of the deficits. Importantly, in the group receiving human EG, no side effects on behaviour were observed throughout the study; interestingly, these animals exhibited improvement of sensorimotor function.

### Exogenous human EG sustain brain tissue regeneration

Histological analysis using Nissl staining was used to appreciate and quantify the new tissue in the damaged brain area. In a recent paper, we developed a method for quantifying this tissue[24]. *Figure 4* shows the characterization of the new tissue in the four groups of rats; the dotted black line corresponds to the lesion and the dotted red line identifies the new tissue. Daily injection of conditioned medium for one month did not promote the formation of new tissue (7% *vs*. 15% in medium group). In contrast, intranasal administration of human EG induced a remarkable tissue regeneration, as shown in *Figure 4B, B’ and E*. Strikingly, the quantification of the new tissue reached 24% for human EG *vs*. 10% for vehicle group (*Figure 4E, P=0.016*). An interesting finding was the fact that, when we looked at each animal in detail, we found that in the human EG group one rat with small lesion size had a high percentage of reconstructed tissue (51.0%, *Supplementary* Fig. 2A, B *and E*). This was associated with important rescue of sensorimotor deficits, especially for the asymmetry test (*Supplementary* Fig. 2F-H). We compared this rat with one having similar lesion size but receiving the vehicle. As expected, we found lower tissue regeneration (10.5%, *Supplementary* Fig. 2C-E), with negligible restoration of sensorimotor deficits (*Supplementary* Fig. 2F-H). This result suggests a link between the extent of tissue regeneration and rescue of sensorimotor impairment.

**Figure 4.**
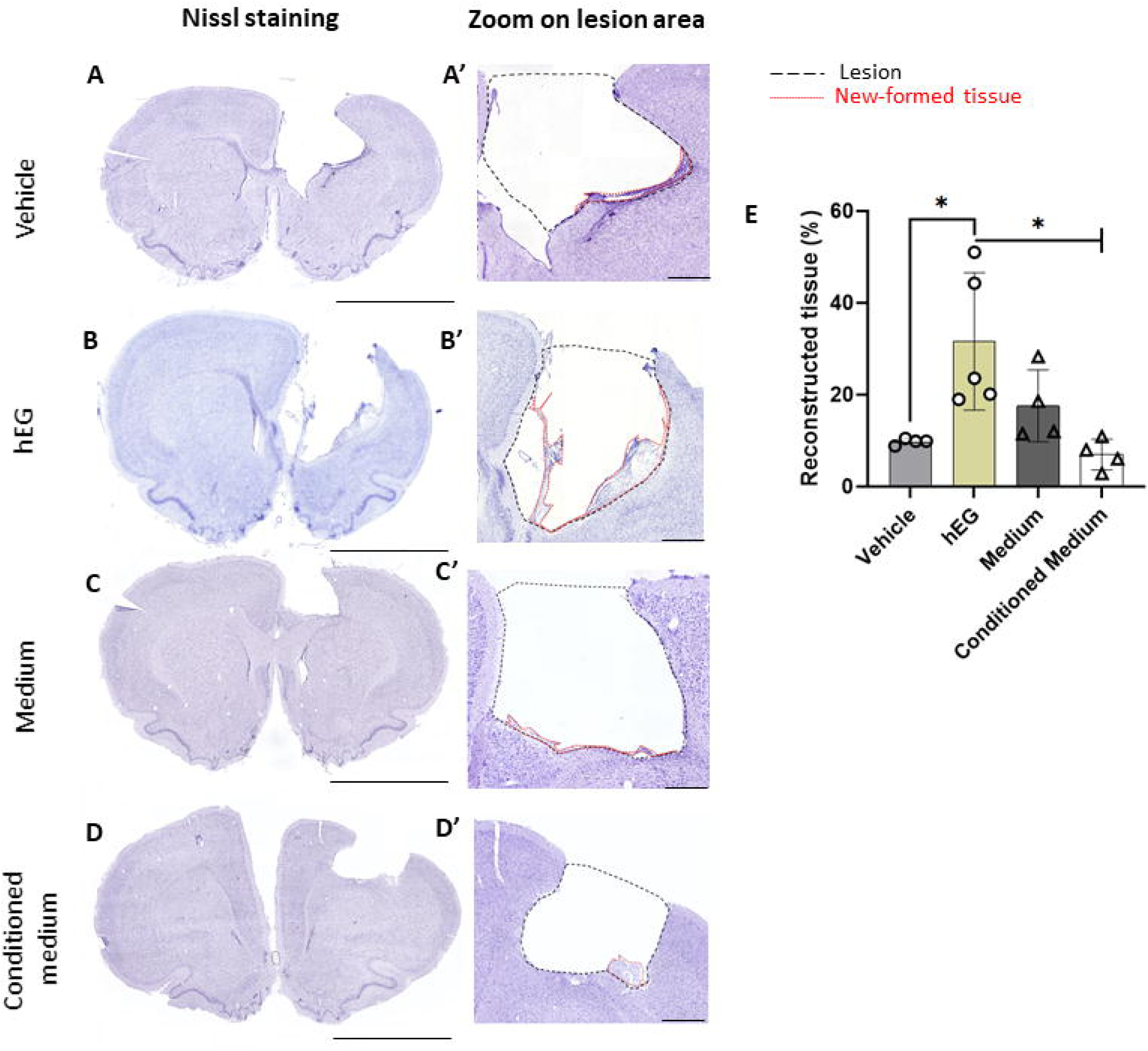
New tissue quantification and characterization. **(A)** Nissl staining on coronal section in rat receiving vehicle. **(A’)** Magnification showing the edges of the lesion (black dashed lines) and the new tissue (red dotted lines) 24 weeks post-injury in vehicle group. **(B)** Nissl staining in rat receiving hEG. **(B’)** Magnification showing the edges of the injury and the new tissue 24 weeks post-injury in hEG group. **(C)** Nissl staining in rat receiving medium. **(C’)** Magnification showing the edges of the lesion and the new tissue 24 weeks post-injury in medium group. **(D)** Nissl staining in rat receiving conditioned medium. **(D’)** Magnification showing the edges of the injury and the new tissue 24 weeks post-injury in conditioned medium group. **(E)** Graph showing the quantification of th new tissue normalized by the lesion volume in the different groups. The percentage of new tissue was significantly increased in rats receiving hEG (*P=0.016* vs vehicle group, and *P=0.016* vs conditioned medium group, Mann Whitney test). The median and the interquartile range are represented. Vehicle group: n=4; hEG group: n=5; Medium group n=4; Conditioned medium group n=4. A, B, C, D scale bars: 5 mm; A’, B’, C’, D’ scale bars: 500 µm.

### Transplantation of human EG induce host tissue remodelling

Markers of inflammation were quantified to evaluate the response to the four experimental conditions. Intranasal administration of human EG had no effect on astrocytes (Gfap^+^, *Figure 5A-F*) and microglia (Iba1^+^, *Figure A, G-K*) infiltration, compared with the other three groups. The staining with Gfap also allowed measuring the thickness of the glial barrier at the edge of the lesion. The glial barrier is a hallmark after brain injury and is mainly formed of astrocytes. Although we did not find a statistically significant difference in barrier thickness in the four groups, we observed a different architecture of this in rats receiving human EG (*Figure 5L-P*), in which the astrocyte barrier appeared loose and less dense, with palisading astrocytes (*Figure 5M*). These results indicate that human EG can induce the remodelling of the injured tissue by reducing the typical tissue hallmarks of brain injury.

**Figure 5.**
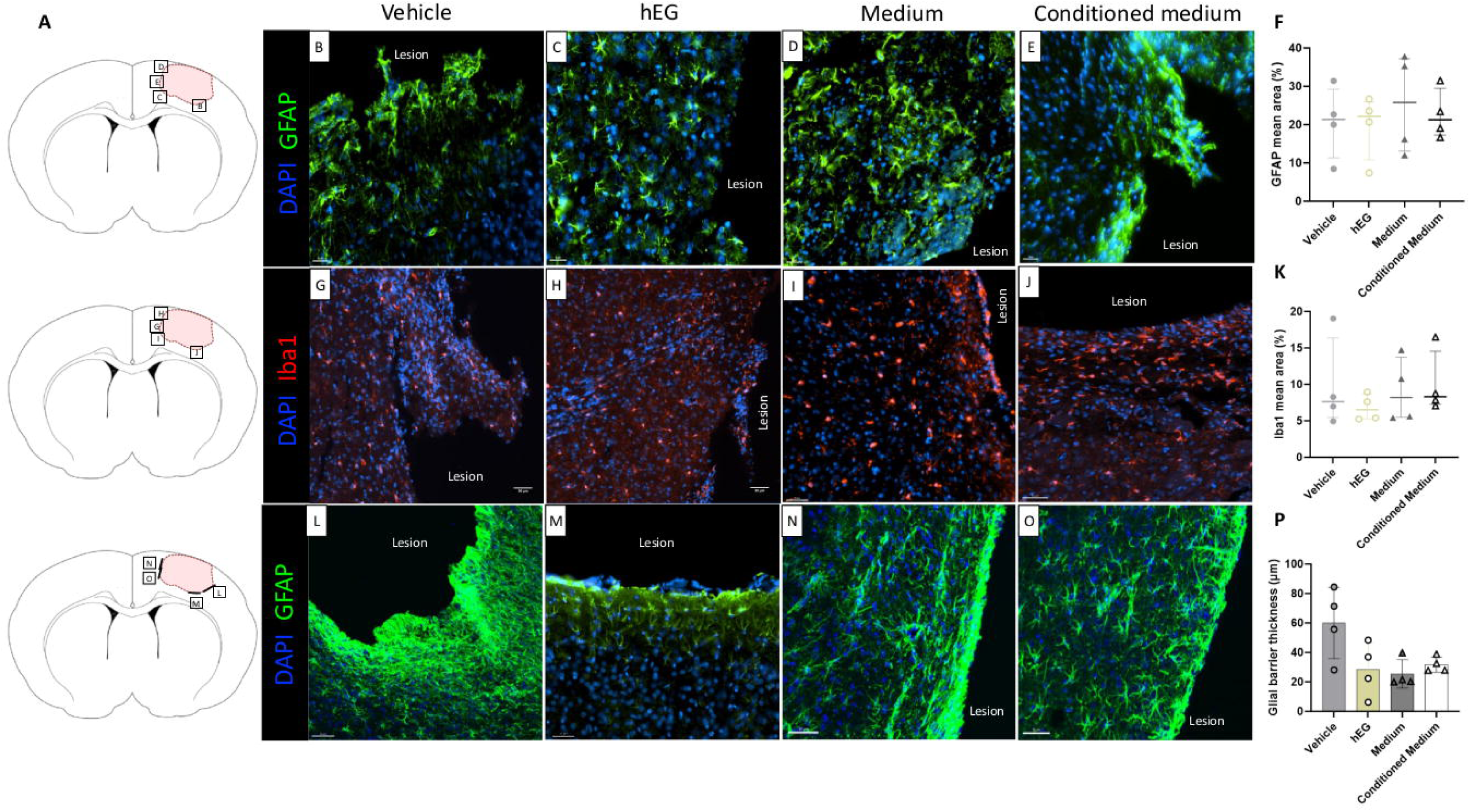
Characterization of the inflammation in the new tissue and at the edge of the lesion 36 weeks post-injury. **(A)** Location of illustrated areas in B-O. **(B-F)** Representative images and quantification of GFAP (to identify astrocytes) in the new tissue. **(G-K)** Representative images and quantification of Iba1 (to identify microglia) in the new tissue. There was no significant difference in the number of immune cells between the groups, as reported in the graphs. **(L-P)** Representative images and quantification of the astroglial barrier (expressed in μm) based on GFAP mean intensity (fluorescence) in the different groups. There was no significant difference between the groups. All graphs show the distribution of individual values and the median ± the interquartile range. For all groups, n=4. Scale bars B-E: 30 μm; G-O: 50 μm. GFAP: glial fibrillary acidic protein.

Angiogenesis is another important criterion of tissue remodelling. Indeed, neovascularization is a positive criterion for the evolution of lesions, which tends more towards regeneration than scar formation. We assessed the extent of neovessels using the endothelial cell marker Lectin. The quantification of vessel surface revealed enhanced neovascularization in the human EG group compared with the other groups (2.3×10^4^ µm^2^*, P=0.029*, *Figure 6A-F*). Interestingly, in the human EG group, the neovessels were found extending into the new tissue (*Figure 6C*).

**Figure 6.**
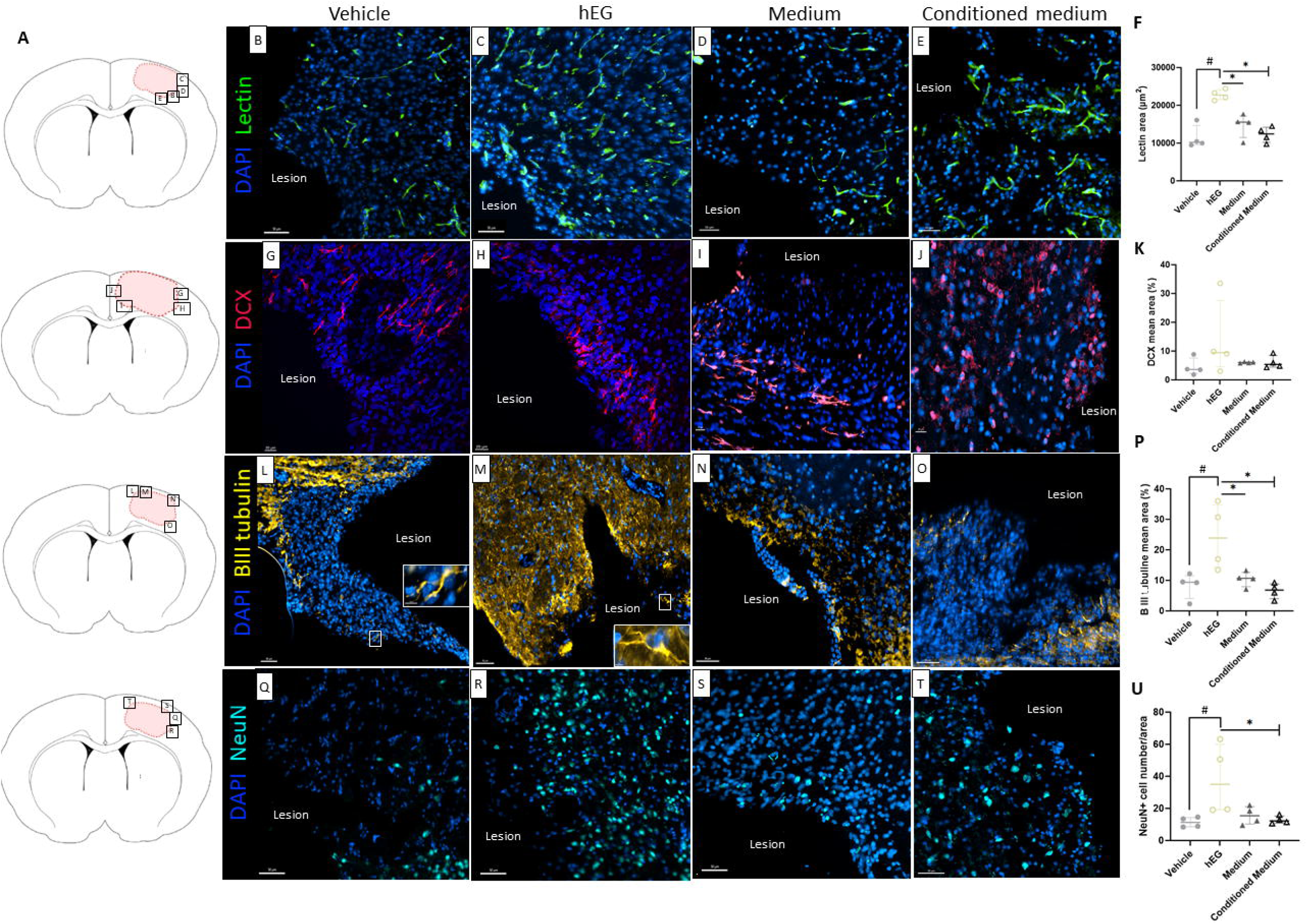
Characterization of the endogenous angiogenesis and neurogenesis in the new tissue 36 weeks post-transplantation. **(A)** Location of illustrated areas in B-T. **(B-F)** Representative images and quantification of Lectin (identify blood vessels) in the new tissue. Positive surface of Lectin is increased in rats receiving hEG compared to the other groups *P=0.029*. **(G-K)** Representative images and quantification of Doublecortin (identify progenitor neurons) in the new tissue. **(L-P)** Representative images and quantification of ßIII tubulin (identify immature neurons) in the new tissue. The inserts show positives ßIII tubulin cell (white arrowheads). A significant difference in the percentage of ßIII Tubulin between vehicle and hEG group was observed (*P=0.029*), also between conditioned medium and hEG group (*P=0.029*), and between medium and hEG group (*P=0.029*). **(Q-U)** Representative images and quantification of NeuN (to identify mature neurons) in the new tissue. A significant difference of NeuN- positive cells between the vehicle and hEG group was observed (*P=0.029*), also between the conditioned medium and hEG group (*P=0.029*). Mann-Whitney tests were performed. All graphs show the distribution of individual values and the median ± the interquartile range. For all groups, n=4. Scale bars G-J: 20 μm; B-E and L-T: 50 µm. DCX: Doublecortin, NeuN: neuronal nuclei.

Further, we evaluated endogenous neurogenesis in the host tissue using three markers corresponding to the different stages of neuronal differentiation: doublecortin (DCX) for neuronal progenitors, betaIII tubulin (βIIItub) for immature neurons, and NeuN for mature neurons. In all groups, we observed the presence of Dcx^+^ cells in the new tissue, indicating the migration of neuronal progenitors from the subventricular zone (SVZ, neurogenic niche) *(Figure 6G-K)*. This suggests that, in our brain injury model, the neurogenesis process was still present thirty-six weeks post-injury. Outstandingly, intranasal administration of human EG induced an increase in both mature (NeuN: 35% *vs*. 11% in vehicle group, *P=0.029*, *Figure 6Q-U*) and immature neurons (βIIItub: 24% *vs*. 9% in vehicle group, *P=0.029*, *Figure 6L-P*), in the new tissue. These results indicate that exogenous human EG importantly contributed to tissue remodelling and enhanced endogenous angiogenesis and neurogenesis.

### Intranasally administered human EG migrate to the lesion area, but do not over proliferate

Following the observation that human EG administration had a beneficial impact on the evolution of the injured tissue, in the subsequent phase of the study we focused on rats receiving human EG and compared them with the vehicle group.

The staining with the human specific nuclei marker, shown in red in *Figure 7*, enabled the identification, localization, and quantification of human transplanted cells after intranasal administration (*Figure 7A-E*). Intriguingly, cells positives for human nuclei marker were identified thirty-six weeks post-injury, which demonstrates their long-term survival. This is an important requisite for cell-based therapy. Quantification revealed a density gradient-like distribution (*Figure 7F*) of human cells in the host brain, with a higher number of cells in the injured site (ipsilesional cortex) compared to the healthy hemisphere (contalesional cortex) (*Figure 7D and E*, respectively). Further, higher numbers of human nuclei-positive cells were identified at the lesion site corresponding to bregma level 2 to 0 slices (median: 4597.5 human nuclei^+^ cells/slice, *Figure 7F*). Interestingly, these cells did not remain in the olfactory bulb or in the frontal region (data not shown), nor were they detected in the region caudally to the lesion site (*Figure 7F*). Together, these findings indicate that transplanted human EG migrated, survived, and integrated into the host brain, preferentially but not exclusively, in the injured site. By reason of the long duration of the study, we wanted to verify the absence of tumorigenic effect after the administration of exogenous human EG. Thus, we evaluated the expression of classic markers of proliferation. Ki67-MIB1 corresponds to the proliferation marker specific to human cells, allowing us to determine whether exogenous human EG were proliferating into the host brain. No Ki67-MIB1^+^ human cells were observed in brain sections (*Figure 7I-I’*). This is an important finding, as it underlines the safety of human EG (xenograft). In parallel, we evaluated the proliferation rate of host (rat) cells, by using Ki67-B56, which labels cycling cells. We found Ki67-B56^+^ cells in the two groups of rats (*Figure 7G-G’ and H-H’*) and we quantified them at the edge of the ventricles, which served as a control of proliferative area (*green arrow*), and in the new tissue (*dark arrow*). Along the ventricles the percentage of cells was comparable in the two groups (24.5% in vehicle group, 21.2% in human EG group, *Figure 7J*); however increased numbers of proliferating cells were evident in the new tissue for the group receiving human EG (12.4% *vs*. 8.8% in vehicle group, *Figure 7J).* The findings demonstrate that tissue remodelling is still ongoing thirty-six weeks post-injury; however, no evidence of tumorigenic risk related to the administration of human EG was found.

**Figure 7.**
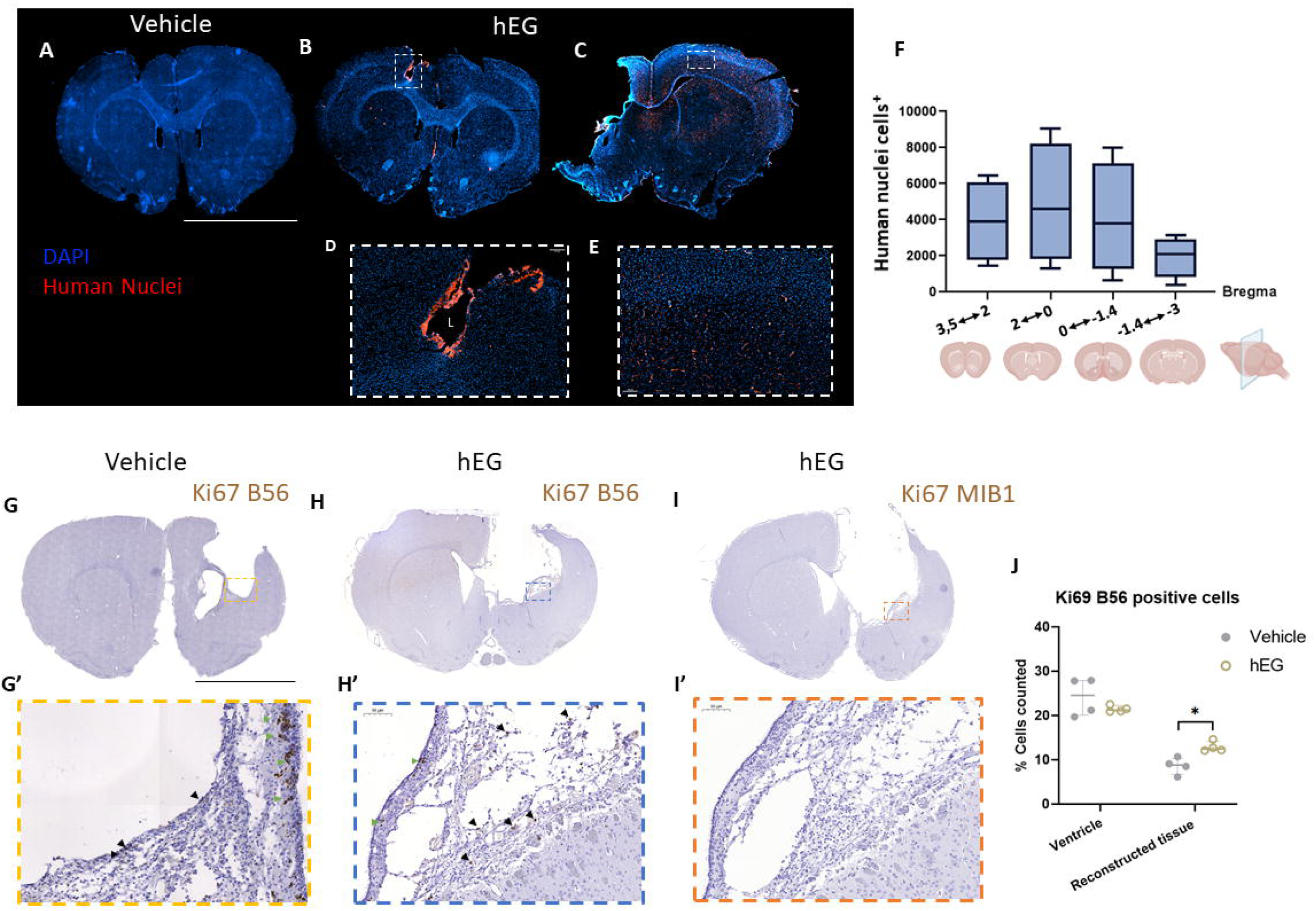
Identification, location, and quantification of hEG 36 weeks post-transplantation. **(A)** Coronal brain section of rat receiving vehicle. Negative control for Human nuclei antibody. **(B-E)** Coronal brain section of rats receiving hEG. Representative images showing human cell location. **(F)** Quantification of hEG in brain sections from 3.5mm bregma to −3mm bregma. Data for each notch box are represented as median ± interquartile range of twelve slices obtained from n=4 rats. A total of 48 sections were quantified. Proliferation rate of human and rat cells. **(G-H)** Representative images of Ki67-B56 staining for vehicle group (G) and hEG group (H). Magnification reveals positive host cells in proliferation (brown). **(I)** Representative image of Ki67-MIB1 staining for the hEG group, showing no human cells positive for the proliferation marker. **(J)** Ki67 positive cell quantification in selected regions of interest with representative images from ventricle (green arrows) and new tissue (dark arrows). Data are represented as individual values with median ± interquartile range. The % of positive Ki67-B56 cells was increased in the new tissue in rats receiving hEG (Mann Whitney test *P=0.028*). Ki67-B56: Rat specific marker of proliferation; Ki67-MIB1: Human specific marker of proliferation. Scale bars (A-C, G-I): 5 mm; scale bars (D, E): 150 μm; scale bars (G’-I’): 50 µm, L=lesion.

### Transplanted human EG generate neurons that integrate with the host brain tissue

To determine the fate of human EG following their arrival in the injured brain, we used the STEM121 antibody, which specifically labels the cytoplasm of human cells. Compared to human nuclei staining, STEM121 allowed to appreciate the morphology of positive cells. We observed that STEM121^+^ cells had a neuronal-like body, as shown in the ipsilesional area (*Figure 8A and B, left*). To confirm our hypothesis, we double-stained brain slices with the neuronal specific marker NeuN, which labels mature neurons, and quantified them. Impressively, we detected 95% of double STEM121^+^/NeuN^+^ cells (*Figure 8B and E*, middle and right, white arrowheads). As depicted in *Figure 8B* (*yellow arrowhead*), some STEM121^+^ neurons showed the morphology of an interneuron. Double STEM121^+^/NeuN^+^ cells were detected as well in the contralesional area (*data not shown*). In parallel, we used glial specific markers GFAP and S100 to evaluate whether human EG retained their glial nature once in the injured brain. We found that only a few cells (5%) were double STEM121^+^/S100^+^ or STEM121^+^/GFAP^+^ (*Figure 8E and Supplementary* Fig. 3).

**Figure 8.**
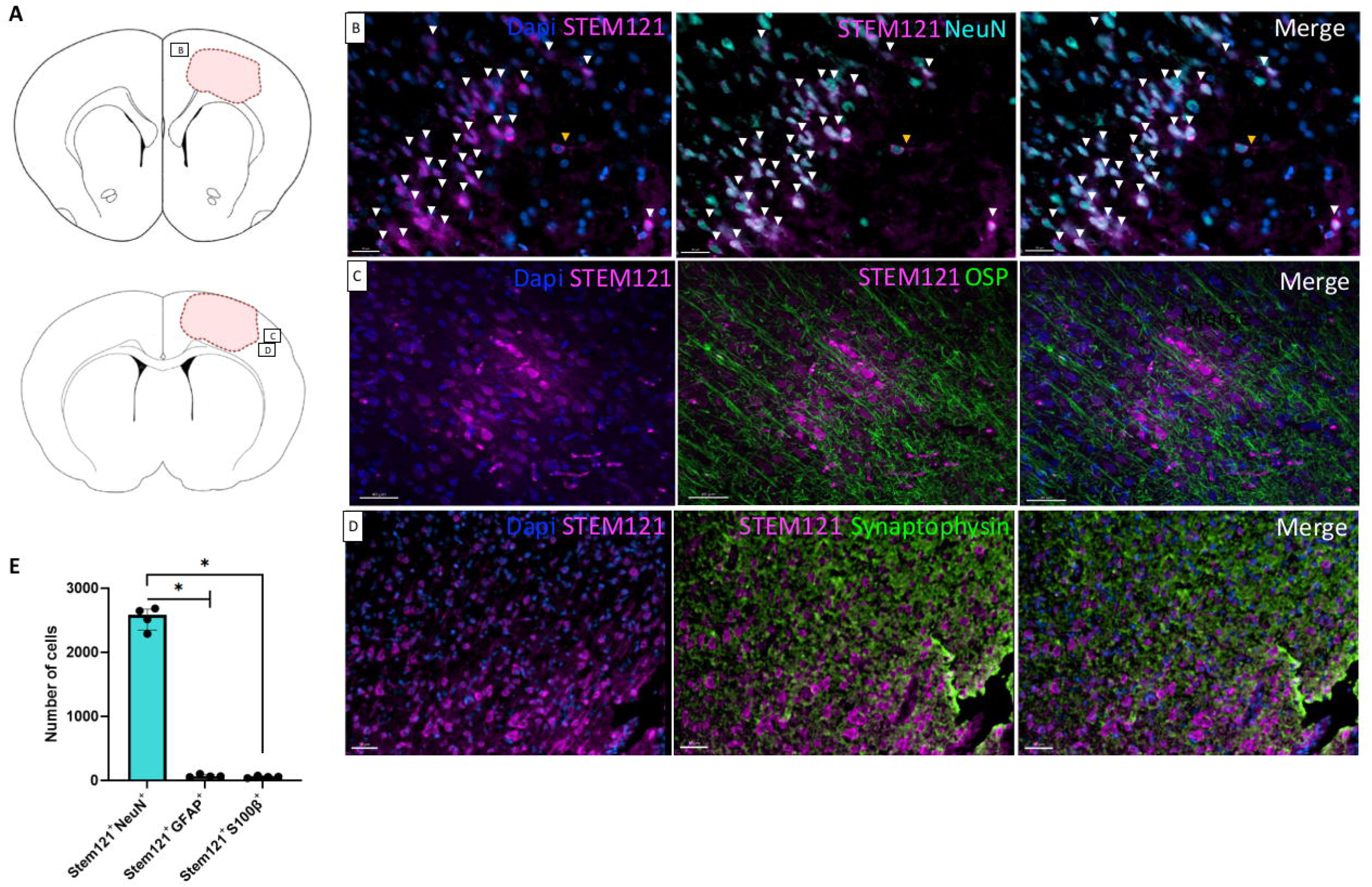
hEG generated neurons anatomically integrated into the injured brain. **(A)** Location of illustrated areas. **(B)** Representative images of hEG stained with human cytoplasmic marker STEM121 and neuronal nuclei NeuN (double-positive: white arrow). The yellow arrow indicates a double-positive cell with an interneuron morphology. Left: Dapi and STEM121 staining, middle: STEM121 and NeuN staining, right: merge. **(C)** Representative images of intranasal hEG stained with human cytoplasmic marker STEM121, and oligodendrocytes with Osp marker specific for myelin. Left: Dapi and STEM121 staining, middle: STEM121 and Osp staining, right: merge. **(D)** Representative images of hEG stained with human cytoplasmic marker STEM121 and Synaptophysin, marker for presynaptic vesicles in neurons. Left: Dapi and STEM121 staining, middle: STEM121 and Synaptophysin staining, right: merge. **(E)** Graph showing the number of engrafted cells double positive for STEM121/NeuN (mature neurons) or STEM121/GFAP or STEM121/S100β (glial markers) (*P=0.029*, Mann Whitney test). Individual values and the median ± the interquartile range are represented. Four counting frames (645 x 628 μm) for two sections per rat were quantified for n=4 receiving hEG. A total of 32 fields were quantified.

To evaluate if neurons generated by human EG successfully integrated with the host brain, we used two specific markers: the Osp (oligodendrocyte-specific protein) marker for myelin and the synaptophysin marker for synapses. Excitingly, we found that STEM121^+^ neurons were integrated and aligned in a neat pattern into the brain cortex and were surrounded by host oligodendrocytes (Osp^+^ cells), indicating that they were myelinated (*Figure 8C*). The reconstruction of the 3D image (*Movie V1*) allows for the clear observation of the alignment and support provided by oligodendrocytes. It depicts two distinct zones within the injured hemisphere: one corresponding to the newly formed tissue and the other situated at a distance from the injury site. Additionally, we found positive synaptophysin staining around STEM121^+^ neurons, which indicated the formation of synapses, a marker of neuronal connectivity *(Figure 8D)*. These findings show the robust integration of EG-derived neurons with the host tissue.

## Discussion

This is the first study to demonstrate the safety and regenerative potential of human EG for the treatment of brain injury. Notably, we show that human EG migrated, survived, integrated with the rat brain tissue, and became neurons. These neurons were myelinated and established synaptic connections with the host tissue. Furthermore, human EG enhanced tissue remodelling, endogenous neurogenesis, and angiogenesis. Together, these effects resulted in marked tissue regeneration.

The evaluation of the safety and efficacy of cell therapies in valuable animal models is an essential prerequisite for the translation into clinical trials. In this study, we used a rat model of brain injury that induces long-lasting sensorimotor deficits, allowing us to follow clinical signs and ultimately measure functional improvement, or deterioration, in injured animals. Two unprecedented aspects of our study are the fact that our rats were immunocompetent and the long duration of the protocol, up to thirty-six weeks. This strategy allowed us to assess that there were no side effects associated with human EG xenotransplantation. Specifically, the evaluation of clinical signs (observation and behavioral tests) and brain tissue anatomy (MRI in live animals and *post-mortem* histology in brain slices) showed no evidence of toxicity in rats receiving human EG.

In our model of long-lasting deficits, we observed a marked improvement in sensorimotor function, particularly in the grip test, between eight and twenty-four weeks in rats receiving human EG. However, no significant differences were appreciable for the other behavioural tests. The integration of transplanted cells with the host tissue is a gradual process that may require considerable time to establish functional connections and, ultimately, functional recovery after brain injury. It would be beneficial for future studies to extend the observation period to evaluate the long-term impact of human EG on functional recovery.

Given the human origin of the donor cells used in our study, assessing the inflammation around the injury and in the new tissue was essential. The evaluation of glial barrier[53] and microglial infiltration[54–56], showed that, compared to the other groups, human EG administration did not exacerbate these signs. In support of the safety of adult human EG, the utilization of a human specific cell marker, Ki67-MIB1, did not evince any indications of human cell proliferation thirty-six weeks following their administration. Conversely, we observed proliferation of host cells (Ki67-B56^+^) in the group receiving human EG. This proliferation was observed in the newly formed tissue, but not in other regions, indicating that the transplantation of human cells resulted in the regeneration of host tissue without the formation of aberrant lesions.

Tissue remodelling is necessary for effective tissue regeneration after brain injury[57]. Interestingly, in our study, the organization of astrocytes forming the glial barrier, was found to be permissive in rats that received human EG, in comparison to the other groups. This was due to the orientation of astrocytes, which appeared palisading. Together, these findings confirmed that the administration of human EG significantly contributed to tissue remodelling.

In this context, and to establish the therapeutic potential of human EG in our model of brain damage, it was essential to quantify the new tissue. Interestingly, we found 24% of new tissue following the administration of human EG, in comparison to 10% in the vehicle group. This percentage represents a remarkable degree of regeneration in relation to the lesion volume. To date, no other studies have quantified the reconstructed tissue following cell transplantation in the injured brain, preventing a comparison of our finding with other results.

The potential for quantifying new tissue also permitted the correlation of this parameter with other variables, such as the recovery of sensorimotor deficit. Interestingly, in the human EG group, the highest percentage of new tissue (51%) was found in a rat with a small lesion size, and this regeneration correlated with significant functional recovery compared to a rat in the vehicle group with a comparable lesion size. The detailed analysis indicated a notable recovery of motor function, particularly evident in the asymmetry test. This is intriguing, as the asymmetry test focuses on shoulder movement, thus revealing early motor improvements[58,59]. The observed functional improvement can be attributed to the presence of the new tissue. Given the limited size of the injury in this rat, we hypothesize that the dose of human EG (1×10 cells) was effective to induce substantial tissue regeneration, which was accompanied by quantifiable functional recovery. Most studies testing the feasibility of intranasal delivered cells in brain injury models have used about one million cells[46,49,60,61]. For example, Vanessa Donega’s study in young mice tested different doses of mesenchymal stem cells (MSCs) ten days after injury[62]. They found that doses of 0.5×10 and 1×10 MSCs improved sensorimotor function, but a lower dose of 0.25×10 did not[62]. We selected a dose (1×10 human EG) that was consistent with the aforementioned studies. Modifying the number of transplanted cells in accordance with lesion size might have a greater effect on functional recovery. However, this hypothesis needs to be confirmed using different doses of cells and the implementation of longer-term behavioural assessments.

The increase in newly formed tissue observed in the human EG group could be due to increased host cell proliferation, as evidenced by positive staining for the specific marker Ki67-B56, discussed above. This proliferation was characterized by enhanced endogenous angiogenesis (lectin^+^ cells) and neurogenesis (βIIITub^+^ and NeuN^+^ cells). The combined effects of increased cell proliferation, angiogenesis and neurogenesis induced by human EG collectively contributed to the remarkable tissue regeneration observed in our study.

A noteworthy finding of our study is the migratory capacity of human EG, which makes the intranasal pathway particularly advantageous to the transplantation of these cells. Human nuclei^+^ cells were detected mainly at the lesion site, and across brain sections (rostral to caudal), exhibiting a gradient-like distribution. The quantification indicated a median value of human nuclei^+^ cells per slice of 3 000, with a total of 20 000 positive cells counted in eight slices per rat, representing a percentage of 18.8% (median value) of intranasally delivered human EG. This result is remarkable in the context of cell-based strategies. Indeed, one of the limitations of other cell sources, such as stem or progenitor cells, is their clearance after transplantation. Only two articles identified in the literature made a quantification after local graft of neural stem/progenitor cells[63], or ENS-derived progenitor cells[64] in mice with irradiation-induced brain injury. The first one found 3.6% of the total number of injected cells five months after cell transplantation[63], while the second one identified only 0.1% of donor cells sixteen weeks after transplantation[64]. In our study, human EG were administered intranasally, necessitating the cells to migrate to the injured region of the brain, in contrast to the aforementioned studies which used local injection, which makes our finding notable.

Noteworthily, within the injured brain, we identified human EG-derived mature neurons, STEM121^+^ and NeuN^+^ cells, which accounted for 95% of total STEM121^+^ cells. This was the most notable outcome of our study, clearly demonstrating the regenerative potential of human EG *in vivo*. In contrast, we found that only 5% of human cells retained their glial nature (GFAP^+^ or S100^+^) in the injured brain. Interestingly, no double positive βIIITub/STEM121 or Dcx/STEM121 cells were detected, suggesting that, thirty-six weeks post-injury, exogenous human EG did not exhibit a neuronal progenitor/immature nature. These *in vivo* data were supported by the *in vitro* evidence that primary mature human EG expressed the ASCL1 transcription factor, indicating their neurogenic potential, without the need of reprogramming. This is an important advantage, compared to other cell sources used for regenerative purposes[64–67]. To date, the neurogenic potential of ENS glia has been described in the mouse intestine, where they may undergo neuronal differentiation under specific conditions[52,68]. Our study is the first to provide evidence that exogenous human EG give rise to neurons in the injured brain. Further investigations at different time points would help to understand the fate of human EG after intranasal administration, when exactly they become neurons and what subtype (excitatory or inhibitory) they are.

Finally, mature neurons generated by human EG exhibited a high degree of integration into the brain cortex, revealing alignment within the host tissue. Furthermore, surrounding host oligodendrocytes myelinated the axons of the neurons, and positive synaptophysin staining indicated that these neurons possessed the requisite components for establishing functional connections. However, we could not directly assess whether these neurons were functional. This type of investigation requires the use of additional animals and imaging techniques enabling real-time detection of labelled human cells in living animals. We are currently exploring this point in a longitudinal study following the fate of human EG at different time points (earlier and later than thirty-six weeks), and using MRI and PET techniques to mechanistically understand the neurorepair potential of these cells after acute brain injury.

## Conclusions

Overall, our study demonstrates that human EG have a therapeutic potential and can be safely used for repair strategies for acute brain injury. This is the first study of its kind and has the potential to be a significant turning point in the field of regenerative medicine.

### Limitations

In this study, we used a brain injury model (malonate injection) known to induce secondary excitotoxic lesions that mimic ischaemic stroke. Anatomically, malonate injection causes massive tissue damage and cell loss. This allowed us to assess the regenerative effect of human EG transplantation in an obvious pathological situation. However, more appropriate models of cerebral ischaemia are needed to bring our approach closer to the clinical situation.

## Declarations

### Ethics approval and consent to participate

The protocol was approved by the “Direction Départementale de la Protection des Populations *de la Haute – Garonne*” and the “Comité d’éthique pour l’expérimentation animale Midi*-Pyrénées*” (protocol n° APAFIS#22419 2019101115259327v5, approved in 2019).

The collection of human gut samples (n=3) was subject to an ethical protocol approved by the national application for the management of the conservation of elements of the human body (CODECOH, protocol: DC - 2015-2443). Informed consent was obtained from all subjects involved in the study.

### Data availability statement

Data will be made available on reasonable request to the corresponding author.

### Competing Interests

The authors report no competing interests.

### Funding

N.C. is supported by the Foundation “Gueules Cassées” (Grant numbers 24-2021, 22-2022, 11- 2023, 18-2024). We thank the Foundation “Gueules Cassées” (Grant numbers 17-2024 to CC; 55-2021 to IL) and the National Research Agency (ANR) (Grant number ANR-19-ASTR-0027 to CC and IL) for financial support.

### Author’s contributions

N.C.: conceptualization, experiments, data analysis, writing original manuscript; E.R.: experiments, data analysis; F.D.: MRI acquisition and analysis, editing; M.C. and M.P.: experiments; L.R.: technical help; E.B. and B.B.: obtaining informed consent, surgical resection of human intestinal tissue; N.V.: responsible for human sample collection, editing; I.R.L.: experiments, data analysis, editing; I.L.: funding acquisition, editing; C.C.: conceptualization, experiments, data analysis, supervision, funding acquisition, writing original manuscript.

## Acknowledgements

We thank Dr Costanza Simoncini for precious help in MRI data analysis, and Mr David Sagnat for technical assistance with human samples. We thank the Non-Invasive Exploration Service, the Experimental Histopathology Platform, and the Animal Experiment Facilities of the US006/CREFRE-Anexplo, Inserm/UT3/ENVT; the Image Core Facility of the Infinity Institute (Inserm) headed by Sophie Allart. We are grateful to Prof Hideki Enomoto and Dr Mukhamad Sunardi (Kobe University, Japan), Dr Vassilis Pachnis and Dr Anna Laddach (The Crick Institute, London, UK), and Dr Alice Le Friec (Toulouse University) for their valuable advice.

The authors declare that they have not used AI-generated work in this manuscript.

## List of Abbreviations

ENS: enteric nervous system
EG: enteric glia
PFA: paraformaldehyde
FCS: foetal calf serum
GFAP: glia fibrillary acidic protein
SOX10: SRY-Box Transcription Factor 10
α-SMA: alpha smooth muscle actin
NeuN: NEUronal Nuclei
ASCL1: Anti-Achaete-scute homolog 1
NSS: neurological severity scale
DCX: doublecortin
βIIItub: betaIII tubulin
SVZ: subventricular zone
Osp: oligodendrocyte-specific protein
MSCs: mesenchymal stem cells

## Notes

### Competing Interest Statement

The authors have declared no competing interest.

### Summary of Updates

Manuscript content updated: supplemental material has been integrated into the main manuscript. Figures have been revised and their order changed.

