## Supplemental Figures for "Long-term effects of xenotransplantation of human enteric glia in immunocompetent rats with brain injury"

**Supplementary Figures**

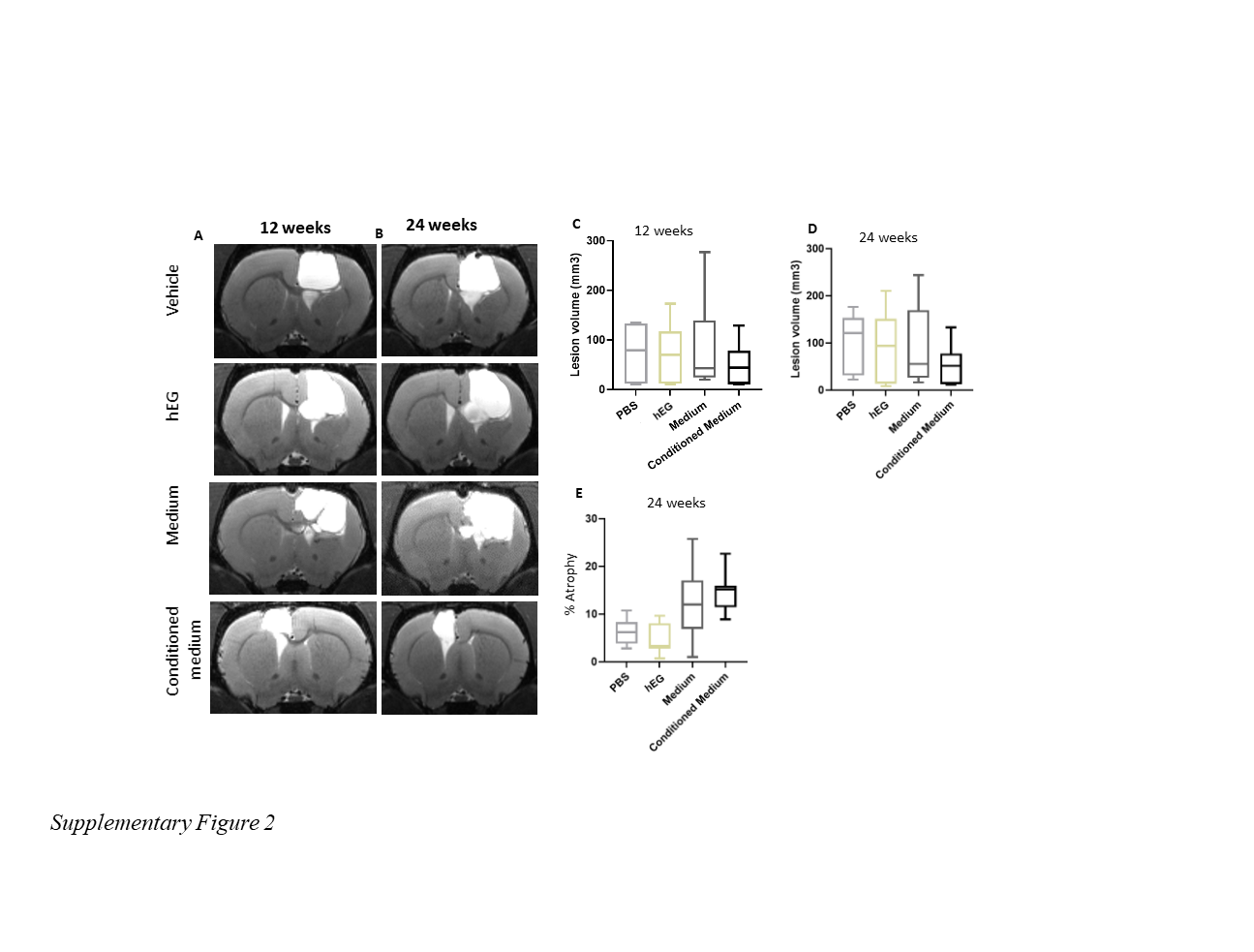
***Supplementary Figure 1****:* MRI follow-up 12 weeks and 24 weeks post-injury. (A) Brain T2 MRI images from rats at 12 weeks are shown: Vehicle, hEG, medium, and conditioned medium. (B) Brain T2 MRI images from rats at 24 weeks are shown: Vehicle, hEG, medium, and conditioned medium. (C) Notch box plot showing the distribution of lesion volumes, expressed in mm^3^, 12 weeks after the lesion. (D) Notch box plot showing the distribution of lesion volumes, expressed in mm^3^, 24 weeks after the lesion. The lesion volume corresponds to lesion volume + injured ventricle volume – healthy ventricle volume + atrophy. (E) Notch box plot showing the distribution of atrophy in %, 24 weeks after the lesion. Vehicle group: *n=7*; hEG group: *n=7*; medium group: *n=7*; conditioned medium group: *n=7*. MRI: magnetic resonance imaging.

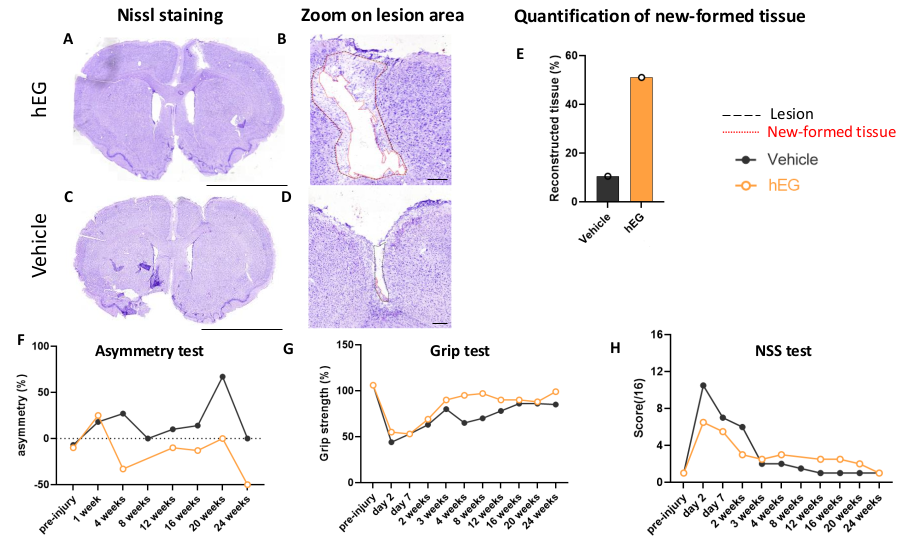

***Supplementary Figure 2****:* *New tissue and behavior tests in hEG vs. Vehicle treatment*. (A) Nissl staining on section in a rat receiving hEG. This section corresponds to the rat with a small lesion in the hEG group, which showed a remarkable new tissue. (B) Magnification showing the edges of the lesion (black dashed lines) and the new tissue (red dotted lines) 36 weeks post-injury. (C) Nissl staining on section in a rat receiving vehicle. This section corresponds to the rat with a small lesion in the vehicle group. (D) Magnification showing the edges of the lesion and the new tissue 36 weeks post-injury. (E) Graph showing the quantification of the new tissue. The rat receiving hEG showed 51% of new tissue vs 10.5% for rat receiving vehicle. (F) The limb-use asymmetry test measures the asymmetric use of the limb in %. (G) The grip strength test shows the grip strength of contralateral paw expressed as %. (H) The neurologic scale score measures sensory-motor impairments (score out of 16). The orange curves correspond to the rat receiving hEG, while the black curve correspond to the rat receiving vehicle. Scale bars: A, C: 5 mm; B, D: 500 µm, Vehicle group: *n=1*; hEG group *n=1*.

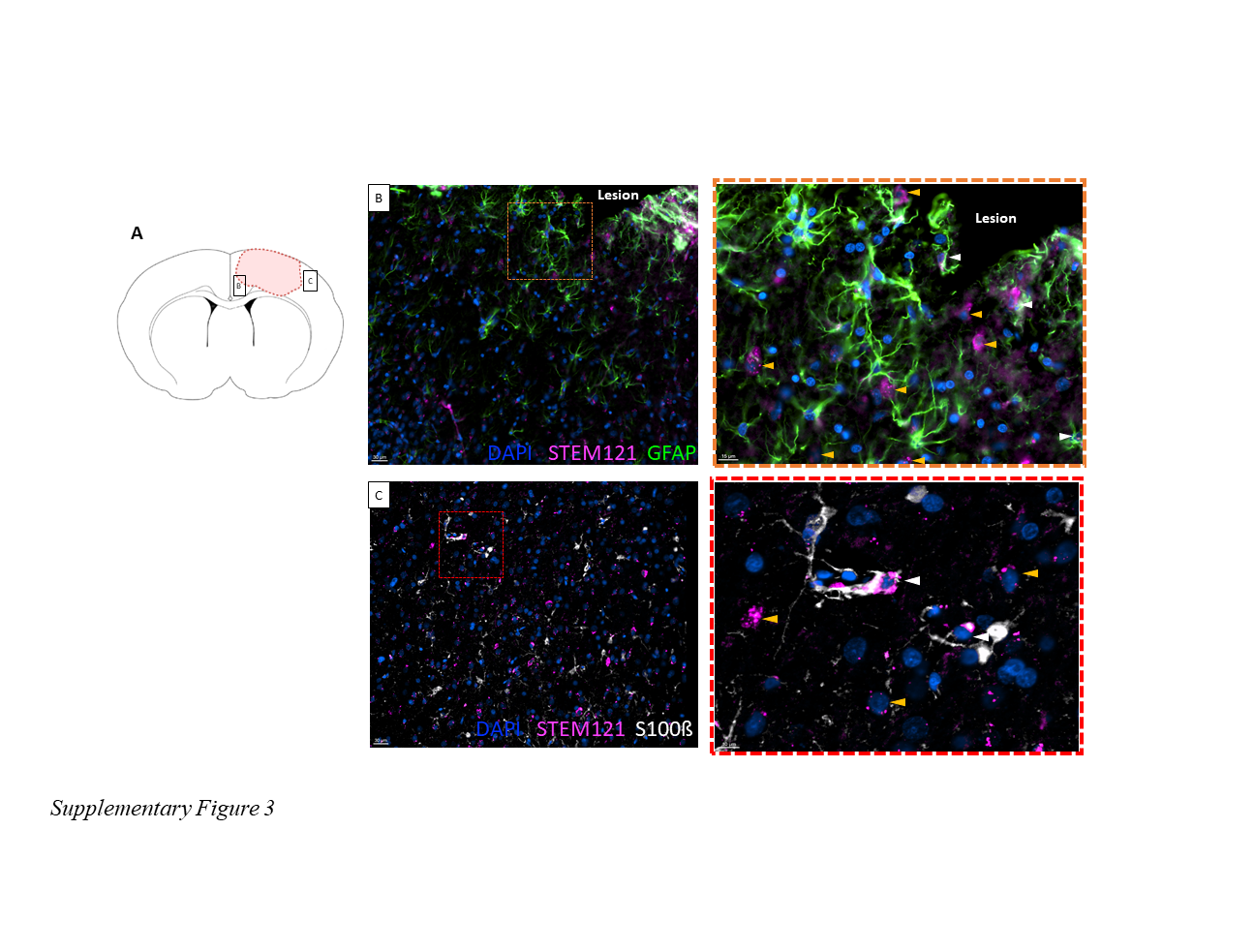
***Supplementary Figure 3****:* Staining of glial markers in hEG after intranasal delivery. (A) Location of illustrated areas. (B) Representative image showing human cytoplasmic marker STEM121 (magenta) and GFAP (green) staining. The crosshatched orange square shows magnification of double positive cell (white arrow) and STEM121^+^/GFAP^-^ cells (yellow arrow). Scale bars=50 μm in the original image and 15 μm in the enlarged image. (C) Representative image showing STEM121 (magenta) and S100ß (white) staining. The crosshatched red square shows magnification of double positive cells (white arrow) and STEM121^+^/S100ß^-^ cells (yellow arrow). Only a few double positive cells are found around the lesion area. Scale bars=50 μm in the original image and 10 μm in the enlarged image.
